# Comparative genomics of *Sympodiorosea* identifies genome evolution mediated through selective pressure on the metabolic gene repertoire

**DOI:** 10.1101/2025.11.25.690603

**Authors:** Jacoby C. Robinson, Aileen Berasategui, Quimi Vidaurre Montoya, Andre Rodrigues, Zoe Zimmerman, Yuliana Christoper, Hermógenes Fernández-Marín, Timothy D. Read, Nicole M. Gerardo

## Abstract

Biological interactions involving host-associated fungi are manifestations of chemistry shaped over evolutionary time. For antagonistic fungi, specialization to host fungi is often facilitated through the acquisition of genes encoding novel secretions, including proteins and specialized metabolites that shape interactions with host defenses, as well the acquisition of nutrients. Alternatively, novel function may arise from diversifying selection eliciting evolutionary innovation in inert secretions. Through specialization, fungal metabolic capacity can serve as an imprint of the selective forces imposed on the antagonist and provide insights into its natural history and occupied niche. Here, we conduct a phylogenomic investigation of antagonistic ascomycetes in the genus *Sympodiorosea*, which are commonly found in basal fungus-growing ant gardens. *Sympodiorosea* and closely related genera of *Escovopsis, Luteomyces, and Escovopsioides* are canonically referenced as virulent mycoparasites, however, recent work has illuminated the possibility for diverse, non-virulent species to emerge within these attine-associated fungi. We explored genomic variation in *Sympodiorosea* to gain insight into the genomic potential for alternative lifestyles, focusing on diversity and evolution of metabolic genes. Our study revealed a constrained selective landscape across the *Sympodiorosea* genome. However, outcomes of *in vitro* interactions with host fungi are diverse and predictable based on the antagonists’ ant-species-of-origin, suggesting functional diversification. Phylogenomics reveals that gain/loss events of genes involved in secretions (secretome) are potential drivers of diversification. Additionally, we demonstrate that purifying selection acts more intensely on secretome-related genes relative to other genes. In contrast, few genes from the secretome experience diversifying selection, suggesting a mechanistic role for driving both functional differences between strains and host-specialization. Comparative genomics including other fungi within the family *Hypocreales* reveals that *Sympodiorosea* has experienced expansions and contractions in proteases genes that are discordant from expectations under a strictly mycoparasitic lifestyle, indicating either the potential for an alternative lifestyle within ant gardens, or that *Sympodiorosea* recently evolved from other niches. These results provide novel insight into genomic evolution of these fungi and inform future experimental studies of the ecology and chemistry of interactions within the complex fungus-growing ant symbiosis.

## Introduction

Fungal parasites and antagonists vary tremendously in terms of their host range and the modes used to identify, contact, and infect hosts. Much of their evolutionary success is reliant on their capacity to synthesize and secrete a diverse array of chemical compounds that mediate interactions with their hosts and with competitors (Case et al., 2022; de Bekker et al., 2017; Keller, 2019). While the ecological, economic and epidemiological importance of some fungal metabolites have been elucidated, less is known about how metabolic diversity across fungi evolves. Our gap in understanding is magnified by the difficulty of culturing many fungal species (Meyer et al., 2016) and the polynucleated genomic structure of some fungi, which makes them less amenable to genomic and computational approaches than other microbes (Kooij et al., 2015; Meyer et al., 2016). Furthermore, a multi-pronged approach that combines experimental and comparative approaches is needed to generate a mechanistic model of genome evolution through the lens of the metabolic gene repertoire an organism contains. Here, to gain insight into metabolic evolution of antagonistic fungi, we used a natural system with an ancient history of association that is well-suited for investigation of how metabolic genes evolve (Berasategui et al., 2024; de Man et al., 2016; Gotting et al., 2022). Ascomycete fungi in the genus *Sympodiorosea* (*Hypocreales*: *Hypocreaceae*) have only ever been found in association with gardens of fungus-growing Attine ants belonging to the ‘lower Attines’ and are considered parasites to the ants’ basidiomycete cultivars (*i.e.,* the fungi that the ants cultivate for food) (Augustin et al., 2017; Bot et al., 2001; Currie, Mueller, et al., 1999; Currie & Stuart, 2001). The cultivars and ants have an array of defenses that can reduce the impacts of fungal antagonists (Currie et al., 2006; Currie, Scott, et al., 1999; Heine et al., 2018; Kyle et al., 2023), likely explaining why *Sympodiorosea* can persist in seemingly healthy gardens in nature. These defenses, as well as interactions with microbial competitors within the gardens, likely have impacted *Sympodiorosea* evolution. With the identification of bioactive metabolites in closely-related fungi (Heine et al., 2018) and recent broad-scale genomic investigations of metabolic potential of *Sympodiorosea* and closely-related fungi associated with fungus-growing ants (Berasategui et al., 2024; Gotting et al., 2022), our primary goals were to assess how metabolic gene diversity within the *Sympodiorosea* genus varies across strains and how selection shapes this diversity within the context of the antagonistic interaction between *Sympodiorosea*, the ants’ cultivars, and, as a consequence, the ants themselves.

The fungus-growing ant system is well-suited to investigate the impact of complex symbiotic interactions on fungal evolution (Currie et al., 2003; Gerardo et al., 2004; Mueller et al., 1998). To date, there are four described genera in the family *Hypocreaceae* of putatively parasitic microfungi associated with fungus-growing ants (Montoya et al., 2021; Osti & Rodrigues, 2018). These include the well described genus *Escovopsis*, *Escovopsiodes* and the newly described genera *Sympodiorosea* and *Luteomyces* (Montoya et al., 2021, 2023), which were split from *Escovopsis*. For these ascomycetes, interactions with the ants’ gardens’ microbial communities are likely mediated by chemistry and secretions produced by the cultivated fungi (de Oliveira et al., 2023; Gerardo et al., 2006; Goes et al., 2024), as well as those produced by other bacteria and fungi commonly found in gardens (Currie et al., 2006; Currie, Scott, et al., 1999). Recent genomic analyses described functional variation in regions related to metabolism across these genera (Berasategui et al., 2024; Gotting et al., 2022), revealing that secondary metabolism broadly differs across these taxa. This suggests that their chemistry is evolving according to genus-specific selective pressures. For a symbiotic fungus, genes encoding metabolic secretions are likely to experience significant selection due to the necessity to produce compounds critical for infection, signaling, defense, and nutrient acquisition (Barrett et al., 2020; Girard et al., 2013; Jia et al., 2023; Saciloto-de-Oliveira et al., 2023; Wang et al., 2022) and due to the energetic costs associated with producing these secretions. This suggests that distributions of secondary metabolites may reveal the history of adaptation to specific niches or hosts, and that signatures of selection on the genes encoding for critical metabolites may be different than those of other genes not evolving in response to dynamic species interactions.

For *Sympodiorosea* spp., evolution may be shaped by interactions with the ants, the fungi cultivated by the ants, and by other microbes in the ants’ gardens. *Sympodiorosea* is commonly isolated from the gardens of a subset of the Attine ants, including ants in the genera *Cyphomyrmex*, *Myrmicocrypta*, and *Mycetophylax* (Custodio & Rodrigues, 2019; Montoya et al., 2021). *Cyphomyrmex* obligately cultivate basidiomycete fungi in both filamentous and yeast forms and are geographically distributed across central America, reaching from southern parts of the United States to the Amazon Basin (Currie et al., 2003; Mueller et al., 1998; Schultz & Brady, 2008). Previous work within the *Cyphomyrmex* association has demonstrated considerable diversity in interaction outcomes between *Sympodiorosea* and cultivar strains (Birnbaum & Gerardo, 2016). Acting as an antagonist to the cultivars, *Sympodiorosea* can recognize and overgrow cultivars grown by the native ant species from whose gardens they were isolated – less often they can overcome defenses of non-native cultivars grown by other ant species (Birnbaum & Gerardo, 2016; Gerardo et al., 2004). These results indicate genus and species-level mechanisms of specialization for the antagonistic fungi. Consistent with these findings, previous work using marker genes revealed that more closely related *Sympodiorosea* are found in *Cyphomyrmex* colonies with more closely related cultivars (Birnbaum & Gerardo, 2016; Gerardo et al., 2004). Specifically, while *C. longiscapus* is closely-related and ecologically similar to *C. muelleri,* the two species cultivate fungi from separate clades (‘lower fungal cultivars Clade 1’ and ‘lower fungal cultivars Clade 2’), whereas *C. costatus* are not found in the same habitats and are more distantly related yet cultivate fungi genetically similar to the cultivars of *C. muelleri* from ‘Clade 2’ (Green et al., 2002; Mehdiabadi et al., 2012; Schultz et al., 2024). Diversification of *Sympodiorosea* has never been studied at the level of the genome nor in relation to genes that may be functionally important for interactions within the context of association with particular cultivars and their ant farmers.

Here, we use comparative genomics to describe the phylogenomic landscape of *Sympodiorosea.* After establishing a whole genome-based phylogeny, we assess diversification across the *Sympodiorosea* genus, specifically focusing on genomic regions related to primary and secondary metabolism. Our results provide support for the naming of the newly described genus *Sympodiorosea*, indicate that variation in metabolic genes is constrained by strong purifying selection and demonstrate that, at the genomic scale, *Sympodiorosea* likely are evolving in response to interactions with both the cultivars and the ants. These results indicate a further need to explore the chemical mechanisms that underpin interactions between associated fungi.

## Materials & Methods

### Assessing morphology and *in vitro* host overgrowth

Twenty-two *Sympodiorosea* and nine cultivars *(i.e.* the fungi cultivated by ants for food) were obtained from a culture collection maintained at Emory University (Supplementary Table S1). The 22 *Sympodiorosea* comprise 12 strains isolated from *Cyphomyrmex longiscapus* colonies, three strains isolated from *C. costatus* colonies, three strains isolated from *C. muelleri* colonies, two strains isolated from novel species of *Cyphomyrmex* attines (*C*. sp. nov.), one strain isolated from a colony of an unknown species of *Cyphomyrmex* (*C.* sp.), and one strain isolated from a *Myrmicocrypta ednaella* colony (Supplementary Table S1).

Samples were revived from 10% glycerol stocks stored at room temperature. All samples were maintained on Potato Dextrose Agar (PDA), stored at 25 °C and transferred once Petri dishes were overgrown, or every 14 days, depending on growth rate. To assess gross *in vitro* morphology, *Sympodiorosea* were grown in monoculture. The cultures were incubated at 25 °C and photographed every 3 days up to the 28-day mark.

To assess *in vitro* antagonism directed at cultivars hosts, we then set up pairwise bioassays of the 22 *Sympodiorosea* strains with nine cultivar strains. The cultivars included six isolated from colonies of ants in the genus *Cyphomyrmex.* These cultivars are all in the family *Agaricaceae* (order *Agaricales*) and represent two main clades of *Leucocoprinus* cultivars grown by Lower Attines (Mueller et al., 1998; Schultz et al., 2024). Specifically, the three strains isolated from *C. longiscapus* colonies are from ‘lower fungal cultivars clade 1’, and the three strains isolated from *C. muelleri* and *C. costatus* colonies are from ‘lower fungal cultivars clade 2’ (Schultz et al., 2024). The remaining three cultivar strains were isolated from *Apterostigma* spp. colonies. These ‘coral fungus cultivars’, in the family *Pterulaceae* (order A*garicales*), are distantly related to the lower fungal cultivars (Schultz et al., 2024). For each combination of cultivars and *Sympodiorosea*, cultivars were plated, in triplicate, ∼5mm from the edge of a 60mm Petri dish containing PDA by transferring a 5mm plug of fresh mycelium on PDA from a Petri dish. They were allowed to grow for seven days at 25 °C. At day 7, *Sympodiorosea* were plated centrally in the same manner. Experimental plates were incubated at 25 °C and photographed every 3 days up to the 28-day mark. Images of *Sympodiorosea* and cultivars interactions were analyzed in ImageJ v1.53 (Schneider et al., 2012). Interaction fidelity was measured through a binary, where ‘1’ indicates that *Sympodiorosea* overgrew the cultivar and ‘0’ indicates that *Sympodiorosea* was unsuccessful in overgrowing the cultivar because 1) the cultivar inhibited *Sympodiorosea* growth at a distance 2) *Sympodiorosea* was inhibited at the point of contact, or 3) *Sympodiorosea* failed to invade the cultivar hyphae in the event contact occurred (Christopher et al., 2021).

### Whole-genome sequencing of *Sympodiorosea*

Genomic DNA extractions were performed for each of the *Sympodiorosea* samples used in bioassay experiments. Strains were first grown on PDA for seven days, at which point a small plug of each strain was transferred to a new PDA plate to ensure monoclonal conditions. After an additional four days, fresh mycelia from each of the strains was gently scraped off the agar using sterilized spatulas. For each strain, 500 µL of fresh mycelia was collected in 1.5 mL Eppendorf tubes containing 500 µL ddH_2_O and centrifuged at 13000 RPM for five minutes. Care was taken to ensure PDA did not dislodge into the mycelial suspension. The supernatant was removed, and 200 µL lysis buffer (recipe in Supplementary Methods and Figures) and 100 µL of microbeads were added. Two hundred microliters phenol:chloroform:isoamyl (25:24:1) was added to the suspension before vortexing for 30 minutes. Two hundred microliters of TE buffer was added to each tube before centrifugation at 13000 RPM for five minutes. The aqueous layer was removed, and each sample underwent a second extraction with 400 µL phenol:chloroform:isoamyl. DNA was precipitated overnight in 100% methanol at 4C, dried, and resuspended in 50 µL ddH_2_O. All samples were sequenced through OmegaBioservices in 2021 using the Illumina NovaSeq 6000 platform. Manufacturer protocols were used to prepare samples for PE 2×150 short-read sequencing using the NovaSeq 6000 SP Reagent Kit (Modi et al., 2021).

### Ultra-Conserved Element sequencing of Cultivars

Cultivar identity was confirmed by amplification and sequencing of ultra conserved elements (UCEs) from pure cultures, which have been previously used to reconstruct the evolutionary history of Leucocoprinus cultivars. Sequencing was conducted at the Smithsonian Museum for Natural History using protocols outlined in Schultz et al. 2024.

### Genome assembly

We used Trimmomatic (v.0.39) to both trim adapter content and filter all reads for quality. High quality reads were assembled *de novo* using spADES v.3.15.5 (Bankevich et al., 2012), with the “--fungus” and “--careful” flags. BUSCO (Benchmarking Universal Single-Copy Orthologs) (Simão et al., 2015) was used to assess assembly quality by searching against the “Ascomycota_odb10” universal single-copy ortholog database. All samples analyzed showed a “completeness” score > 95%, indicating high quality assemblies with low levels of duplication or fragmentation (Supplementary Table S1).

### Functional annotation

Gene prediction and annotation were performed using the Funannotate (v.1.8.13) pipeline. In short, repeats were identified and soft-masked with tantan (Frith, 2011). Soft-masked genomes underwent gene prediction by first aligning proteomes from *Trichoderma* sp., *Cladobotryum* sp., *Hypomyces rosellus* and *H. perniciosus* using DIAMOND/Exonerate. PASA (Do et al., 2024) gene models were used to train/run three gene prediction tools: Augustus (Stanke & Morgenstern, 2005), SNAP (Bromberg & Rost, 2007), and GlimmerHMM (Delcher et al., 1999). The joint predictions were then passed through EvidenceModeler to provide a consensus prediction (Haas et al., 2008).

We used BlastP searches against the Uniport/SwissProt and MEROPS protein databases to generate functional annotations and identify protease and protease inhibitors, respectively (Altschul et al., 1990). Functional annotations fed through InterProScan5 (Jones et al., 2014) produced putative protein domains and sites as well as Gene Ontology (GO) terms with subsequent alignments inferred by searching against the Eggnog orthology database using emapper v3 (Cantalapiedra et al., 2021). Biosynthetic gene cluster predictions were assigned using fungiSMASH v6 (Blin et al., 2021) using all predictive tools under relaxed parameters. Finally, carbohydrate active enzymes and their putative substrates were annotated using the dbCAN3 server (Zheng et al., 2023) with HMMER v3.3 (Finn et al., 2011).

### Phylogenetic reconstruction and genetic distance

The phylogenetic relationship of *Sympodiorosea* and their close relatives was reconstructed using the BUSCO phylogenomics pipeline (McGowan & Fitzpatrick, 2020). The phylogenetic analysis of intraspecific diversity was bolstered by the inclusion of 46 publicly available proteomes (six additional *Sympodiorosea* spp., 12 *Escovopsis* spp., five *Luteomyces* spp., 11 *Trichoderma* spp., five *Fusarium* spp., two *Hypomyces* spp., one *Cordyceps militaris*, and one *Cladobotryum* sp.) (Supplementary Table S2). The busco_phylogenomics pipeline first identified shared, single-copy orthologs using the BUSCO v5 ‘Hypoceales_odb10’ lineage database. Protein sequences were aligned using MUSCLE (Edgar, 2004), and alignments were trimmed with TrimAl (Capella-Gutiérrez et al., 2009). Trimmed alignments were concatenated into a supermatrix, and the alignment was partitioned by ModelFinder, which was allowed to infer the best evolutionary-based model (Kalyaanamoorthy et al., 2017). The partitioned alignment was used as input for IQTree (Nguyen et al., 2015) to generate the phylogeny using the fast bootstrap flag ‘-B 1000’ with *Cladobrotryum* spp., *Hypomyces rosellum* and *H. perniciousus* as outgroups.

To place our sequences within a larger context of antagonists associated with fungus-growing ants, we generated a maximum-likelihood phylogeny using the elongation factor 1 alpha (tef1-alpha) gene sequences NGphylogeny (Lemoine et al., 2019). We included 157 total samples spanning *Escovopsis*, *Luteomyces*, *Escovopsioides*, and *Sympodiorosea* (Supplementary Table S4). Sequences were aligned using MAFFT (Katoh & Standley, 2013) with default settings, and phylogenetic informative sites were selected via BMGE (Criscuolo & Gribaldo, 2010). The phylogeny was estimated with PhyML with Smart Model Selection (Guindon et al., 2010) using default settings, and branch support was based on SH-like aLRT branch support.

Additionally, we tested for phylogenetic signal of the host cultivar and ant associate using the PhyloSignal package in R. In other words, we tested whether ant species of origin correlates with *Sympodiorosea* relatedness and, similarly, whether cultivar host correlates with relatedness between *Sympodiorosea*. A consensus tree was combined with tip data related to the native ant species cultivating the gardens from which our 22 genome-sequenced strains were isolated as well as six publicly available *Sympodiorosea* genomes (Supplemental Table S2).

For the 22 *Sympodiorosea* strains sequenced as part of this work, average nucleotide identity values were calculated using PyANI (v0.2.10) under default settings. We used MUMmer (NUCmer) to align our 22 sequenced genomes. Pairwise sequence alignment was used to calculate the percentage identity between two given samples.

### Comparisons of metabolic gene content

Given that these strains vary in terms of the ant species with which they were associated in nature and their ability to infect (in *sensu lato* as overgrow) cultivar hosts *in vitro*, we investigated differences in the genes underlying substrate use, both of which could influence host use. Gene presence/absence data tables were generated for carbohydrate active enzymes (CAZyme) and proteases. The presence/absence data were used as input for principal component analysis and heatmap visualization. Principal component analyses were carried out with the function ‘prcomp’ available in the base R stats package and visualized with ggplot2 (Wickham, 2016). All heatmaps in our analyses were generated using the R package pheatmap (Raivo Kolde, 2010).

In addition, putative functional annotations of CAZymes and proteases were converted to counts, and, subsequently, we determined a ratio of these for each genome (CAZyme:protease). These ratios were placed in the larger context of alternative fungal lifestyles by scraping data from Ahrendt et. al (2018), Fig. 5 using WebPlotDigitizer v.4.6 (Ahrendt et al., 2018).

For Biosynthetic Gene Clusters (BGCs), we used BiG-SCAPE to generate sequence similarity matrices by grouping Gene Cluster Families (GCFs) based on sequence similarity and homology, with a cutoff score of 0.5. In the matrix, GCF presence is indicated by ‘1’ and absence by ‘0’. GCF presence/absence was visualized with a heatmap using R. Exemplar BGCs which encode known metabolites from the closely-related species *E. weberi* were selected for synteny analysis using Clinker (Gilchrist & Chooi, 2021) to understand how metabolic potential may be shaped at the locus-level. From the putative predictions, we sought to identify known clusters by searching against the MIBIG database v3.1, which is composed of 2,502 secondary metabolite clusters by using the flag ‘mibig’ in the bigscape.py command.

### Linking *in vitro* overgrowth to Genomic Variation

To test for association between variation in overgrowth, metabolic gene content and phylogenetic relatedness, we used the command “vegdist” from the R package “Vegan” (Oksanen et al., 2001) to generate dissimilarity matrices of Euclidean distance based on host-use and ANI, as well as, protease, CAZyme and BGC presence/absence. We used Spearman’s correlation coefficients when using the ANI dissimilarity matrix in Mantel tests of association. Otherwise, tests were conducted using Kendall’s correlation coefficient due to the qualitative nature of presence/absence data.

### Population Genomics and Selection

To understand the role of selection in diversification of *Sympodiorosea*, we calculated the ratio of nonsynonymous mutations (dN) to synonymous mutations (dS) for each ortholog that was previously assigned a Gene Ontology term. Gene ontology enrichment analyses were executed for our 22 sequenced strains using GenBank formatted annotations supplied to ‘Funannotate compare’ with the flag ‘--run_dN/dS estimate’ using the ‘M0’ null model from codeml (Yang, 2007). The M0 model effectively averages the strength of selection across all sites and branches for a particular gene. The resulting values are conservative estimates of selection, because, to detect selection, it would need to act intensely within a few samples or moderately across all samples. Our data composed of highly similar genomes further reduces the likelihood that selection can be detected, as sufficient time must pass for orthologous genes to acquire identifiable differences in their coding sequences. Ultimately, we seek to identify genes that are under intense selection and that may contribute to the coevolutionary relationship between *Sympodiorosea* and the other organisms within the fungus-growing ant symbiosis.

To evaluate signatures of selection across sequenced genomes, we implemented the free branch model as a conservative estimate of selection, where selection is assumed to be consistent across all sites along all branches (0 < theta < 1). Each ortholog in each genome was then assigned a value corresponding to the calculated ratio of dN to dS (hereafter referred to as dN/dS) (Supplementary 5). An arbitrary of cutoff of dN/dS > 0.45 was set as the threshold for elevated selection. This cutoff is in line with other studies of fungal pathogens/parasites that reported dN/dS ratios ranging between 0.1 - 0.4 for core and accessory genes (Cheng et al., 2015; Olarte et al., 2019; Petersen et al., 2023)

We then selected orthologous protein encoding genes identified as the core proteome in our strains with OrthoFinder and Cluster of Orthologous Genes (COG) database. In total, we identified 5,023 core proteins and assigned dN/dS values to each ortholog in each genome. dN/dS values were binned according to COGs, and t-tests were carried out by comparing the dN/dS distributions between COGs related to metabolism and defense to background genes; principally, we were interested in the COGs: ‘G’ = carbohydrate, ‘U’ = intracellular trafficking, secretion, and vesicular transport, ‘V’ = defense, ‘I’ = lipid, and ‘Q’ = secondary metabolism. Together, these COGs were classified as the secretome.

## Results

### Sequencing and Assembly of *Sympodiorosea*

We shotgun sequenced 22 strains using Illumina technology. Genome assemblies were high quality, with an average of 95% completeness under the “Ascomycota_odb10” dataset using the BUSCO (Benchmarking Universal Single-Copy Orthologs) scoring method. Average assembly size was 29.4 Mb – ranging from 26.05 Mb to 35.8 Mb and harboring, on average, 7,724 protein-coding genes. GC% ranged considerably between 40.1% and 48.3%, with an average of 46.4% (Supplementary Table S1).

### Phylogenomics reveals a weak but significant pattern of ant-of-origin association

Pairwise Average Nucleotide Identity (ANI) between the *Sympodiorosea* genomes ranged from 86% – 100% (Fig. S2). Clustering ANI scores based on Euclidean distances revealed seven distinct clusters. Owing to sometimes large nucleotide sequence differences between genomes, we elected to use protein alignments to reconstruct evolutionary relationships. We extracted 4487 single copy orthologs found in all 68 proteomes (22 strains from this study and 44 other fungi from seven genera (Supplementary Table S2). Our supermatrix alignment of 676,661 amino acids produced a Maximum likelihood phylogeny with strong support (> 95%) for most nodes (Fig. 1, Fig. S3). The bootstrapped phylogeny was congruent with a phylogeny with more samples estimated using tef-1 alpha, a marker gene used in fungal phylogenetics (Fig. S4, Supplementary Table S4). As previously demonstrated (Berasategui et al., 2024; Gotting et al., 2022), these reconstructions revealed that attine-associated antagonists are monophyletic and that *Sympodiorosea* spp. strains form a monophyletic clade within the attine associates.

**Figure 1.**
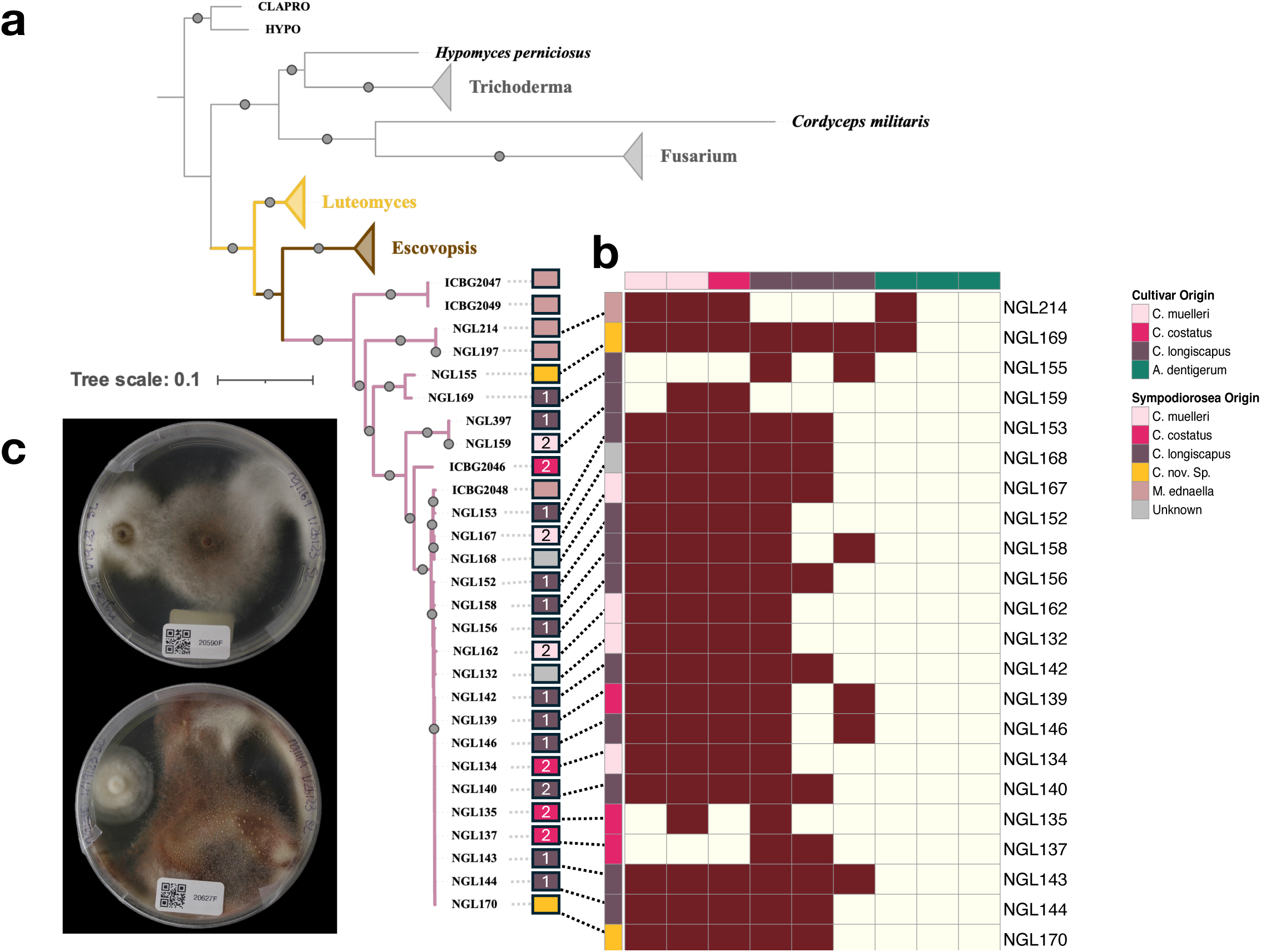
Phylogenomic placement and host-use of whole-genome sequenced *Sympodiorosea*. A) Consensus species tree, based on 4487 shared orthologs, generated from 1000 ultrafast bootstrap replicates. Triangles represent collapsed clades containing outgroups. Non-*Sympodiorosea,* attine-associated symbionts are represented by collapsed clades, with yellow for *Luteomyces* and brown for *Escovopsis*. *Sympodiorosea* strains are indicated on pink branches. A fully resolved phylogeny with all taxa can be found in Fig. S1. Filled black circles indicate ultra-fast bootstrap values of > 90%. The phylogeny indicates that *Sympodiorosea* spp. form a distinct clade sister to *Escovopsis* spp. isolated from higher attine and *Apterostigma* colonies. Colored boxes next to leaves indicate the ant species from which a sample was isolated, with numbers corresponding to associated cultivar clade (*i.e.,* clade 1 or clade 2 in Schultz et al. 2024), when known. Within the *Sympodiorosea,* this study putatively identifies at least six clades that may represent distinct species of *Sympodiorosea.* B) Heatmap of host-use bioassay outcomes. Dashed lines connect position in phylogeny with row in bioassay. Brown filled boxes indicate that the *Sympodiorosea* strain was successful in overgrowing the cultivar, while ivory indicates that *Sympodiorosea* was inhibited by and did not successfully overgrow the cultivar. Filled boxes at the top and to the left indicate the ant-origin of the fungal isolate. Each cell represents the proportion of times a *Sympodiorosea* strain could overgrow a cultivar based on triplicate plates. C) Confrontation assays between *Sympodiorosea* strain NGL169 and a Leucocoprinus Clade 2 cultivar (top) and NGL169 vs. an outgroup cultivar (bottom).

Within *Sympodiorosea*, consistent with the ANI analysis, we the phylogenomics analysis supported at least six well-supported clades (bootstrap values > 95%). Four *Sympodiorosea* isolated from *Myrmicocrypta* colonies, ICBG2047, ICBG2049, NGL197 and NGL214, were basal to those strains isolated from *Cyphomyrmex* colonies. The next most derived clade included NGL169 and NGL155, isolated from colonies of an undescribed *Cyphomyrmex* species and *C. longicapus* respectively. This clade was basal to NGL159, isolated from a *C. muelleri* colony, and NGL397, isolated from a *C. longiscapus* colony. Sister to the NGL159-NGL397 clade was a branch harboring ICBG2046 isolated from a *C. costatus* colony. ICBG2046 was the basal branch for the remaining 19 samples of which only ICBG2048 was non-*Cyphomyrmex*-associated. Branch support was lower within the most derived clade, however, the branches containing NGL153 and ICBG2048 were highly supported (bootstrap = 100), as was an intermediate branch containing NGL168 and NGL167 (bootstrap = 100). Clustered within the largest clade were the remaining 16 *Sympodiorosea* strains: NGL152, NGL158, NGL132, NGL156, NGL140, NGL142, NGL134, NGL137, NGL146, NGL170, NGL144, NGL143, NGL135, NGL139, and a well-supported branch containing NGL162 (bootstrap = 100).

While the seven distinct clades were not each specific to an ant species, assessment of phylogenetic signal at the strain level revealed that the ant-species-of-origin was a phylogenetically conserved trait (Pagel’s λ = 9.42e-01, p < 0.001) (Fig. S5a). Contrastingly, we found no evidence that host cultivar clade was correlated with relatedness between *Sympodiorosea* strains (Pagel’s λ = 6.63e-05, p > 0.05) (Fig. S5b); notably, this trait was known for only 17 of the 22 strains, limiting power to detect an association.

### *Sympodiorosea* preferentially overgrow native hosts

*In vitro* bioassays pairing *Sympodiorosea* with cultivars were performed to assess potential host range. We used cultivars isolated from *C. longiscapus*, *C. muelleri*, and *C. costatus* gardens, as well as cultivars from gardens of ants in the genus *Apterostigma dentigerum* (Supplementary Table S3).

As previously seen (Birnbaum & Gerardo, 2016), we found *Apterostigma-*associated cultivars to be highly resistant to overgrowth by *Sympodiorosea; Sympodiorosea* were only able to overgrow an *Apterostigma*-associated cultivar in 2 of 69 bioassays. In contrast, the same *Sympodiorosea* were able to overgrow *Cyphomyrmex-*associated cultivars in 99 of 138 bioassays.

Within the *Cyphomyrmex* association, we identified clear patterns in overgrowth in relation to cultivar clade (Fig. 1B). *Sympodiorosea* strains isolated from *C. longiscapus* gardens infected *C. longiscapus* cultivars (*Leucocoprinus* clade 1 cultivars) in 64% (27/42) of bioassays. In contrast, *C. longiscapus*-derived *Sympodiorosea* infected *C. muelleri* and *C. costatus* cultivars (*Leucocoprinus* clade 2 cultivars) in 89% (40/45) of bioassays. *C. muelleri*-derived *Sympodiorosea* infected *C. muelleri* and *C. costatus* cultivars (*Leucocoprinus* clade 2 cultivars) in 100% (12/12) of bioassays, but only infected *C. longiscapus* (*Leucocoprinus* clade 1 cultivars) in 42% (5/12) of bioassays. Interestingly, *C. costatus*-derived *Sympodiorosea* infected *C. muelleri* and *C. costatus* cultivars (*Leucocoprinus* clade 2 cultivars) in 44% (4/9) of bioassays, and infected *C. longiscapus* (*Leucocoprinus* clade 1 cultivars) in 56% (5/9) of bioassays. Overall, *Sympodiorosea was* significantly more likely to overgrow their native-type cultivars (*i.e.,* cultivars from the same clade as the *Sympodiorosea*’s garden of origin) than non-native cultivars (Fisher’s exact, p = 0.0015, Supplementary Table S5, Fig. S14). There was a significant correlation between genetic similarity of the strains, based on ANI, and their host range, based on overgrowth patterns. (Mantel test, r = 0.32, p < 0.01).

Our study also captured interactions that diverged from the norm. Strains NGL214, NGL169, NGL155, and NGL159 formed arm-like protrusions composed of mycelial mats that extended from the media, grew along the ‘ceiling’ of Petri dishes, and finally infected the cultivar from the ‘air’ (Fig. S7b). This behavior was specific to interactions with the host; in the absence of hosts, these same strains failed to produce the arm-like protrusions until the plate was nearly covered, at which point protrusions would form. Even when a protrusion-forming strain was initially inhibited by the presence of a cultivar, it could deploy this growth behavior to grow directly into the host while remaining distant from the presumably metabolite-laden media. This trait is specific to the basal lineages of *Sympodiorosea,* suggesting that it may have been lost in the more derived strains.

### Cazymes and Proteases are highly conserved across *Sympodiorosea*

To identify variation in metabolic potential, we surveyed the distributions of gene annotations related to CAZymes, proteases and biosynthetic gene clusters. On average, 228 genes (range 221-232, median = 228) encoding CAZymes were identified across the 28 *Sympodiorosea* strains. CAZyme protein predictions were generally conserved across all strains (Fig. S13a, Supplementary Table S11), with only one gene family, GH18, found to be significantly different in counts across the 22 *Sympodiorosea* genomes sequenced for this study (Fig. S6). GH18 is linked to chitin degradation and was the most abundant CAZyme gene identified in our analysis. Relative proportions of CAZyme classes were conserved, with a per-genome average of 23 protein encoding genes related to ‘auxiliary activities’, five ‘carbohydrate binding module’, 10 ‘carbohydrate esterase’, 134 ‘glycoside hydrolase’, 67 glycosyl transferases’, and five ‘polysaccharide lyase’. From these, we surveyed the putative substrates to gain insight into the lifestyle of *Sympodiorosea*. We found glucan degrading enzymes to be most abundant, with an average of ∼40 protein-encoding genes. Of the glucan enzyme genes, beta-glucan was the most frequently predicted substrate, with an average of 37 genes putatively related to beta-glucan degradation per genome. Glycan degrading enzymes followed, with roughly 30 predictions per genome. An average of 15 chitinases, 14 ligninolytic enzymes, 10 xylanases, five arabinogalactan proteins, five glycogen-degrading enzymes, five pectinases, and four mannoses were predicted per genome.

There was little variation in both individual proteases counts and relative abundances of protease families across the 28 *Sympodiorosea* genomes (Fig S16b). We found serine peptidases most common, with an average of 73 per genome, followed by glutamic peptidases with 67 genes, cysteine peptidases with 55, threonine peptidases with 19, aspartic peptidases with 16, and protease inhibitors with three. While individual counts differed in some protein families across strains, we found no evidence that any proteases are differentially conserved within the *Sympodiorosea*.

The distribution of serine protease families is of particular interest in understanding the lifestyle of a fungus, as these hydrolytic enzymes serve critical functions in nutrient acquisition and virulence in fungi (Figueiredo et al., 2016; Li et al., 2017; Lv et al., 2023; Lysøe et al., 2017; Muszewska et al., 2017). Contractions and expansions of serine protease subfamilies have been suggested as illuminating the lifestyle and niche of a fungus. Within *Sympodiorosea*, we identified a considerable expansion of S9 (mean = 22) and S33 (mean = 21) serine peptidase subfamilies compared to other fungi, which is unexpected for pathogenic fungi in general (Muszewska et al., 2017) (Fig 2b, Supplementary Table S13). Such patterns of expansion are most common in symbiotic fungi of plants (Muszewska et al., 2017). Furthermore, we found contractions of enzymes from the S1 (mean = 4) and S8 (mean = 3) clases and an absence of S15 enzymes, which are commonly expanded in fungal pathogens (Muszewska et al., 2017).

**Figure 2.**
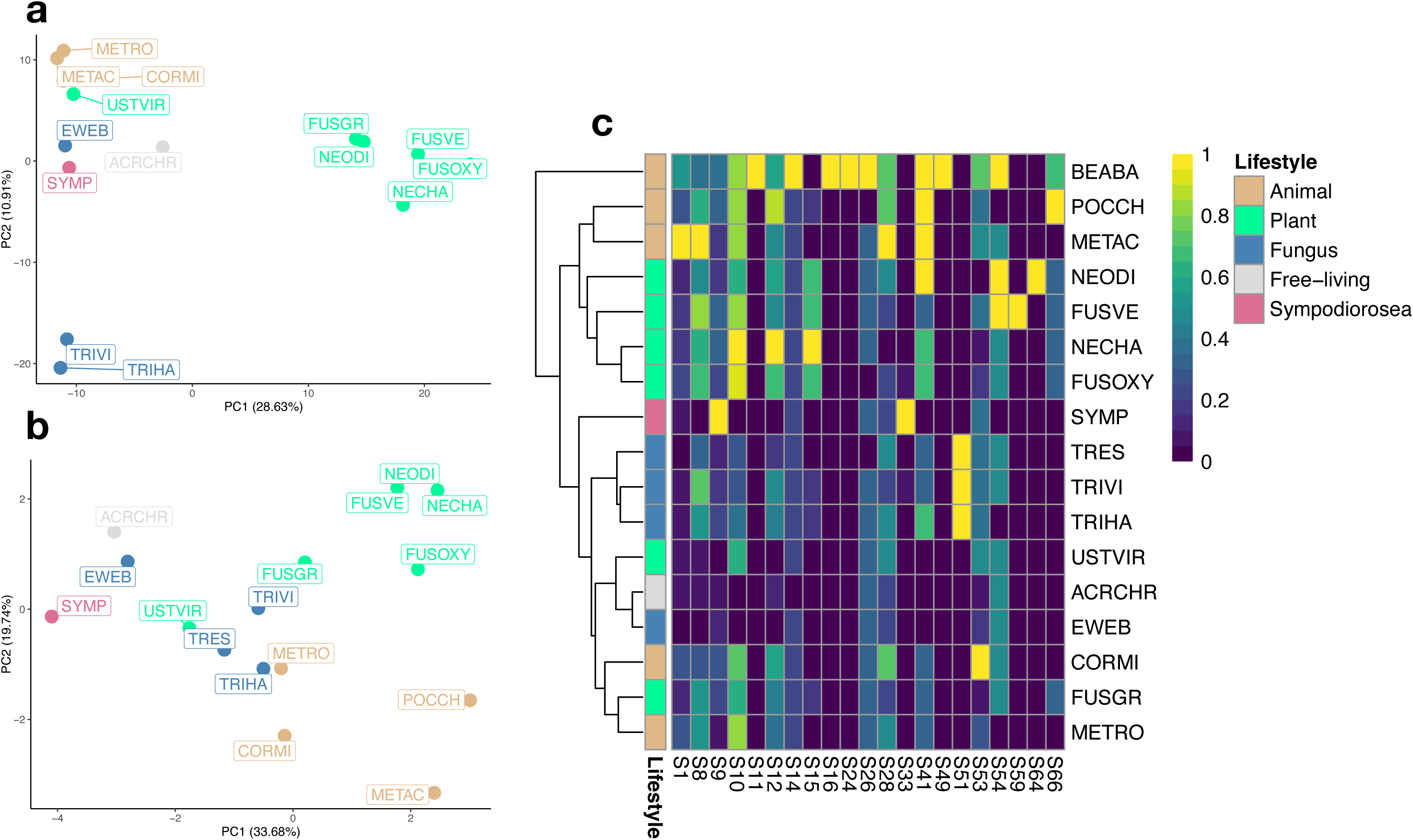
Variation in metabolic potential, focusing on genomic diversity of carbohydrate-active enzymes and proteases. Principal component analyses based on protein annotations related to: A) carbohydrate active enzymes (CAZymes) and B) serine proteases. Colors indicate lifestyle of the sample: animal pathogens = light brown, plant pathogens = light green, Fungal pathogens (mycoparasites) = blue, free-living = grey, Sympodiorosea = pink. C) Proteomes in 2b hierarchically clustered based on genes encoding serine proteases. Columns are normalized to the highest value to enable comparisons between protease families. Lifestyle colors follow the schema of 2a and 2b.

### *Sympodiorosea* are genomically discordant from a strictly mycoparasitic lifestyle

Fungi employ a range of proteases and carbohydrate active enzymes (CAZymes) that are critical for cellular defense and nutrient acquisition respectively. It has been suggested that patterns of contraction and expansion in these gene classes can provide insight into potential lifestyles. Of the 128 CAZymes with an assigned putative substrate, nearly 30% were specific to glucan degradation, with (37/40) enzymes specific to beta-glucan. Twenty three percent were specific to host-glycan degradation/degradation of fungi. Glucan is among the most abundant polysaccharides in fungal tissues, indicating that *Sympodiorosea* have specialized to the most abundant sugar source in fungus-growing ant gardens. Chitin is another polymer abundant in fungal cell walls, and 12% of enzymes found in *Sympodiorosea* were specific to chitin degradation, further supporting our expectation of specialization to the most abundant nutrient source: cultivar-hyphae. Interestingly, *Sympodiorosea* house an equal number of enzymatic genes related to plant degradation (xylan and ligninolytic enzymes) as they do for chitin, a primary component of fungal cell walls.

Within the context of the order *Hypocreales*, our principal component analysis of CAZymes and serine proteases revealed contrasting results regarding how *Sympodiorosea* clusters with other *Hypocreales* fungi (Fig. 2). With regard to CAZyme profiles, we found *Sympodiorosea* to be most similar to its closet relative in the dataset, *E. weberi,* which is a mycoparasite of higher attine agriculture (Reynolds & Currie, 2004). Near to the attine associated fungi were the animal pathogens (two species of *Metarhizium and Cordyceps militaris*) (Fig. 2a). Nearly equidistant from this cluster are two clusters, one containing three *Trichoderma* species and the other containing the *Fusarium*/*Neonectria* species. Obvious clustering breaks down when analyzing the serine protease profiles for the same set of fungi. Notably, attine associated fungi remain clustered, however the mycoparasites from *Trichoderma* replace animal pathogens as the most like attine associates (Fig 2b).

### Biosynthetic Gene Cluster (BGC) composition is indicative of phylogenetic topology

We identified 627 total BGCs grouped into 59 gene cluster families (GCFs), with an average of roughly 22 BGCs per genome (Supplementary Table S9). We identified 59 GCFs in total, 19 of which were shared amongst at least 20 of the genomes; only six GCFs were unique to an individual (Fig. 3). In agreement with recent studies of BGC content in the larger group containing *Escovopsis*, *Luteomyces* and *Sympodiorosea*, we found non-ribosomal peptide synthases to be the most abundant classified BGC, with 21 total identified. Additionally, eight terpene, eight PKS (seven ‘PKS_other’, one PKS Type I) and six hybrid BGCs were found. Twenty-one BGCs were classified as ‘other.’ We identified unique BGCs in NGL214, NGL169, NGL155, and NGL143. The assemblies of NGL155 and NGL143, however, while largely complete, were two of the most fragmented assemblies and as such, the uniqueness of these BGCs should be interpreted cautiously. To understand if genome fragmentation biases BGC predictions, we tested for correlation between the number of predicted BGCs and the number of contigs and found no relationship (r = 0.02).

**Figure 3.**
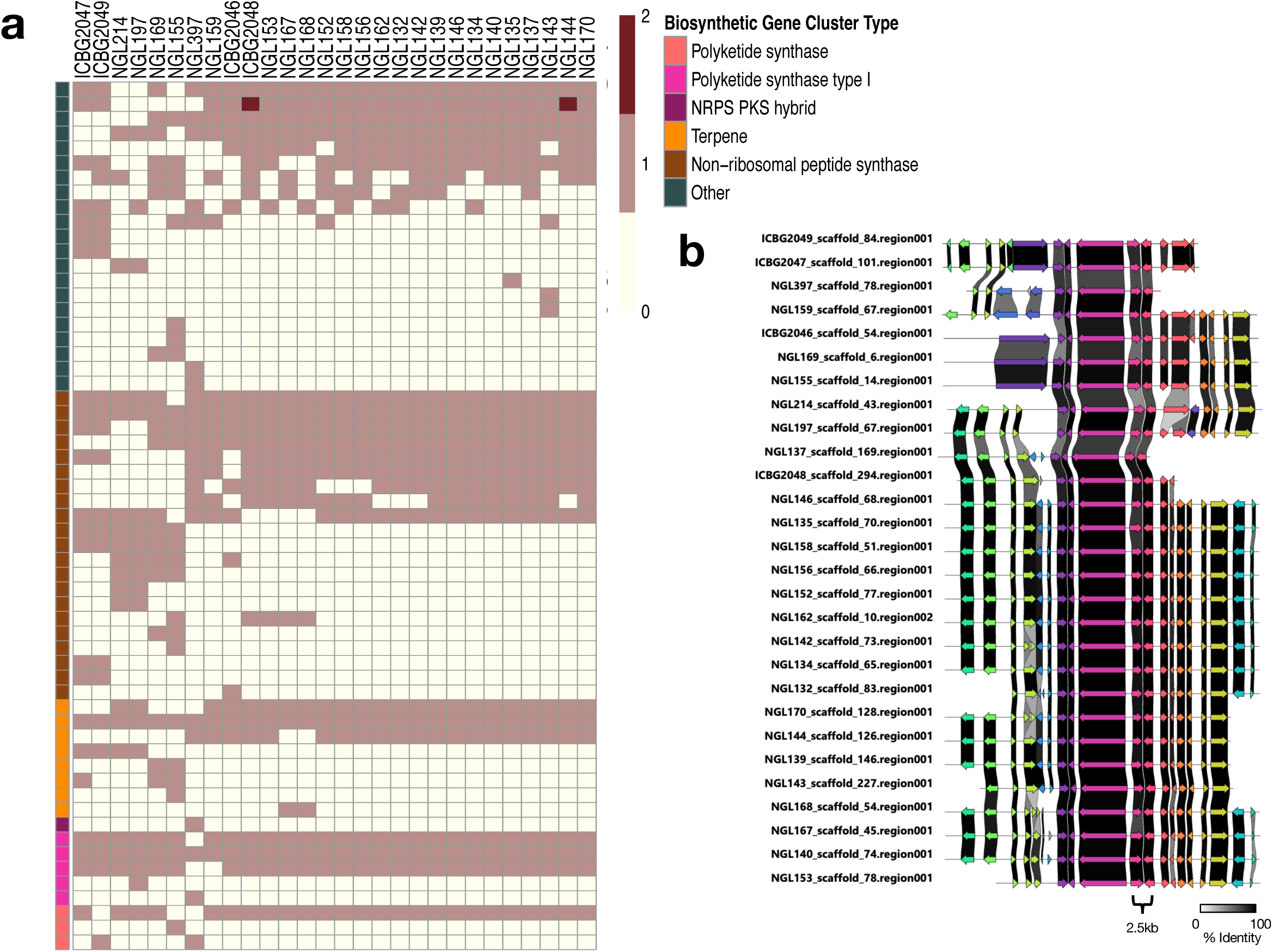
Biosynthetic gene cluster family presence and absence. A) Ivory-colored boxes represent the absence of a Gene Cluster Family (GCF), light brown indicates a single gene copy, and dark brown indicates a duplication of a GCF. Rows, corresponding to *Sympodiorosea* strains, are ordered as in Fig. 1, and columns represent the gene cluster families (GCFs) as identified through BiG-SCAPE. In total, 46 GCFs were identified across the 22 focal strains in our study. Each Biosynthetic gene cluster was assigned to a family based on the ‘Biosynthetic Gene Cluster Type’ annotation column. B) Visualization of an alignment of the BGC likely encoding the compound Melanicidin IV comprised of BGC families FAM00246 and FAM00381.

To test for association between phylogenetic relatedness and biosynthetic gene content, we employed Mantel tests of correlation, with the primary hypothesis that ANI is strongly associated with biosynthetic gene content. Consistent with our expectations, we found that ANI is most strongly associated with the presence/absence of BGCs (r = 0.84, p < 0.0001) followed by CAZymes (r = 0.80, p < 0.0001) and proteases (r = 0.66, p < 0.0005).

### Purifying selection structures the metabolic capacity of *Sympodiorosea*

We hypothesized that clusters of orthologous genes (COGS) involved in metabolic gene traffic experience diversifying selection as a result of chemical evolution in response to the hosts. Similarly, we expected metabolic related COGs to have significantly different distributions of dN/dS values compared to non-metabolic genes, which we hypothesized were more likely to exhibit signatures of strong purifying selection (Supplementary Table S5).

Generally, *Sympodiorosea* exhibit evidence of strong purifying selection (mean dN/dS = 0.134), however, in contrast to our expectations, we found that selection is generally acting to reduce diversity in metabolic-related genes relative to background genes (p < 0.001, Fig. 4). We sought to understand which genes, if any, contributed to the selective differences between metabolic genes and background. Within the secretome, COG ‘Q’, related to secondary metabolism, and COGs G’, ‘I’, and ‘E’ corresponding to carbohydrate and lipid and amino acid metabolism, harbored reduced diversity relative to other metabolic-related COGs. Mean dN/dS values were lower in ‘Q’, ‘G’, ‘I’, and ‘E’ than in the remaining metabolic-COGs, which suggests that these COGs experience strong purifying selection relative to the genome and other non-metabolic related genes.

**Figure 4.**
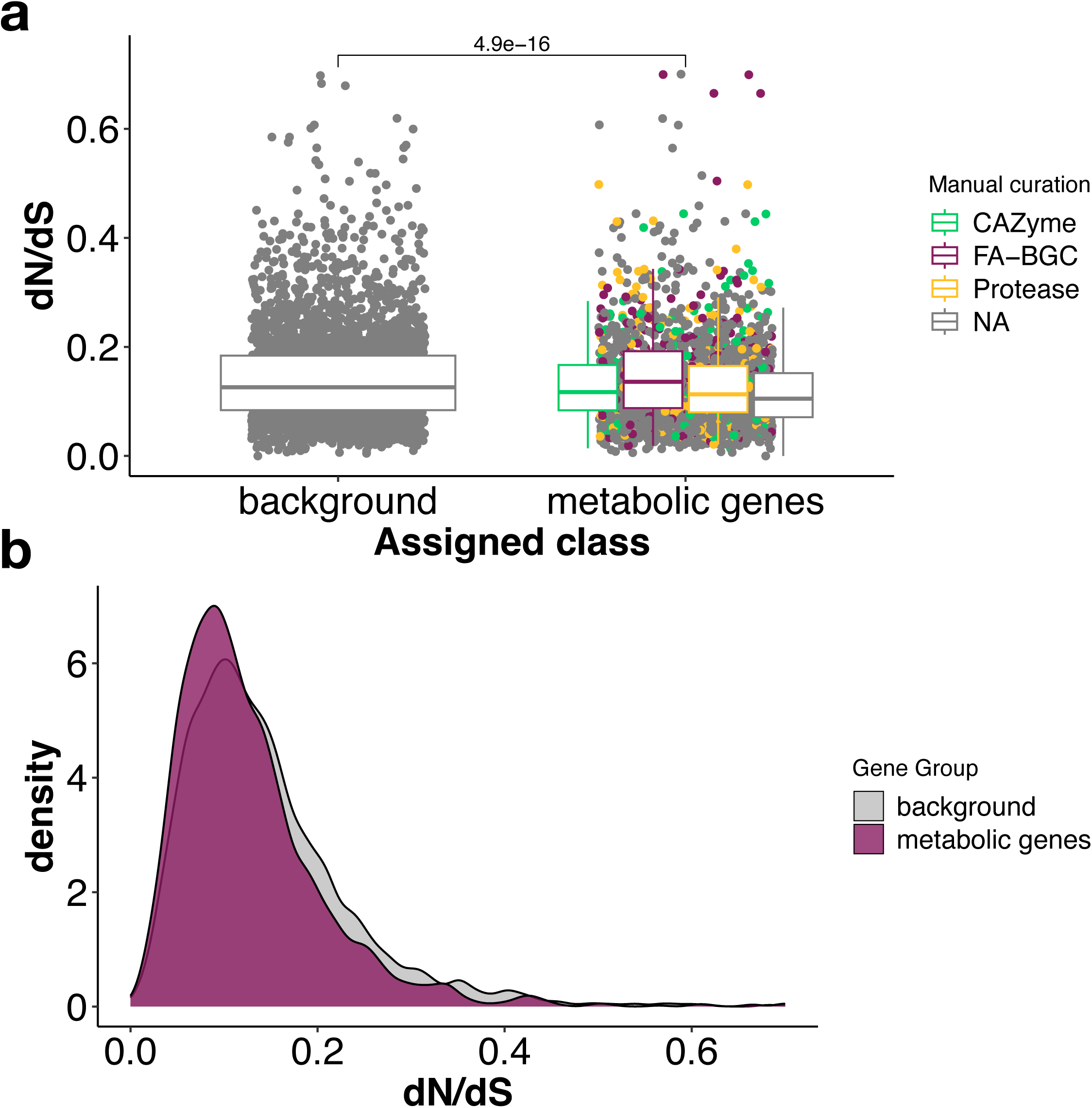
Signatures of Selection. **a)** dN/dS values for 5,049 orthologous protein predictions indicate the associated annotation assigned through manual curation. Metabolism-associated genes in grey were not assigned a gene annotation but were implicated in metabolic activities through gene ontology assignment. Significance indicated at top. **b)** Density plot of dN/dS values colored by their assignment to either the background or metabolic genes.

We sought to confirm these findings by comparing selection acting on genes annotated as proteases, BGCs, and CAZymes to each other and the rest of the genome (Fig. 4). Interestingly, only annotated BGCs were significantly different from the background genomic content (Fig. S12) suggesting a role of diversifying selection in BGC evolution. However, this result is driven in part by a few genes exhibiting signatures of strong diversifying selection; when these outliers are removed, we do not see significant evidence of diversifying selection in BGCs genes relative to the background (p = 0.077, Supplementary Table S7). Twenty-three of the 31 outliers were initially annotated as ‘hypothetical proteins’ without a putative function. Known genes included: PEX29 (Vizeacoumar et al., 2003) responsible for peroxisome size and maintenance, IPO4 (Jäkel et al., 2002) responsible for intracellular trafficking/secretion, MIB1 (Nagase et al., 2000) involved in carbohydrate metabolism, YPD1 (Schruefer et al., 2021) involved in signal transduction, PFY1 (Ostrander & Gorman, 1997) involved in actin polymerization, SUP35 (Dagkesamanskaya & Ter-Avanesyan, 1991) involved in translation termination of GTPase and SQS1 (Kribii et al., 1997) involved in squalene synthesis.

## Discussion

We investigated functional, genomic, and phenotypic diversity of *Sympodiorosea* to gain insight into their evolution in response to interactions within the fungus-growing ant symbiosis. Our study is the first to examine intraspecific diversity at the whole genome-level within this group of fungi. Combined with bioassay experiments, we were able to link genomic variation in metabolism to variation in host interactions. Our study aligns with recent work at a broader taxonomic scale that also demonstrated a clear link between biosynthetic potential, symbiont lifestyle, and phylogenetic relationships (Berasategui et al., 2024; Gotting et al., 2022). These results can be used as the basis for further experiments focusing on the mechanisms driving evolution in this coevolved system.

### Bioassays reveal variation in ability to overgrow host cultivars

Host-specificity may evolve due to prolonged, repeated interactions with a host population. *Sympodiorosea* and its allies have never been isolated outside of an attine garden, suggesting a specialized lifestyle to attine agriculture. Past work has demonstrated fine-level specificity between *Sympodiorosea* and their ‘native hosts’ or the attine species from which they were isolated (Birnbaum & Gerardo, 2016; Custodio & Rodrigues, 2019; Gerardo et al., 2004, 2006; Reynolds & Currie, 2004). Assuming that *Sympodiorosea,* like *Escovopsis* spp., are mycoparasites (REF), successful interactions, from their perspective, occur when the host cultivar has been sensed, initiating chemotaxis towards the host (Gerardo et al., 2006). Then, the *Sympodiorosea* must overcome host-secretions and overgrow the cultivar biomass. For both *Escovopsis* and *Sympodiorosea*, interaction outcomes become predictably incompatible as phylogenetic distance increases for hosts (Birnbaum & Gerardo, 2016; Gerardo et al., 2006). Namely, we confirm findings regarding *in vitro* interactions, where *Sympodiorosea* have preference for native cultivar hosts. Additionally, we confirm previous findings (Birnbaum & Gerardo, 2016) that *C. longiscapus*-associated *Sympodiorosea* overgrow a broader range of hosts than their counterparts from *C. muelleri* and *C. costatus* colonies.

### Phylogenetic relationships of *Sympodiorosea* contextualized

We sequenced 22 *Sympodiorosea* genomes and included six previously sequenced *Sympodiorosea* genomes in our analyses. Of these 28 strains, 22 were isolated from colonies of *Cyphomyrmex* ants. Our results identified a significant phylogenetic signal of native ant host. However, we found no significant phylogenetic signal of cultivar association (*i.e.,* clade 1 or clade 2 cultivars). The lack of phylogenetic signal linked to the native cultivar clade could arise if only a few key genes drive evolutionary interactions with cultivars. For example, cultivar-derived selective pressures may only be detectable within specific gene sets (*e.g.,* the secretome) involved in the maintenance of host interactions. In this case, coevolution between cultivars and *Sympodiorosea* might be masked by selection driven by other forces, including the ants and other microbes, or genetic drift. This however, is not corroborated by previous research using both marker genes and genome-wide amplified length polymorphisms, which both indicated a phylogenetic signal of cultivar association (Birnbaum & Gerardo, 2016; Gerardo et al., 2004). It also contrasts with bioassays, both in our study and in previous studies (Birnbaum & Gerardo, 2016), which demonstrated that *Sympodiorosea* preferentially infect the cultivar type grown by their ‘native’ ant species, indicating specificity. Instead, we hypothesize that small sample sizes collected over long-time scales may mask phylogenetic structure resultant from association with a particular cultivar clade, particularly if evolutionary dynamics include host switching over time.

The explanation is supported by the phylogenetic mixing of *Sympodiorosea* associated with different fungus-growing ant species. *C. muelleri* have been shown to accept the cultivar isolated from *C. longiscapus* colonies, but not vice versa (Mehdiabadi et al., 2012; Mueller et al., 2004), indicating opportunity exists for cultivars, and possibly *Sympodiorosea*, to be transmitted between different species of fungus-growing ant. It is therefore plausible that *Sympodiorosea* coevolves in localized ‘hotspots’, where it is shared between closely-related species. Isolated communities may experience genetic drift, which may appear as specific coevolution within small localities but diffuse over the span of a country. Paired sampling of cultivars and antagonists from the same colonies and at regular time intervals will be necessary to understand the temporal dynamics of specialization.

### Biosynthetic gene clusters are highly conserved

Corroborating other reports, we found that genome reduction has largely eliminated BGC diversity within *Sympodiorosea* relative to other *Hypocreales* fungi, which harbor double the BGCs per genome, on average (Berasategui et al., 2024; Gotting et al., 2022). With an average of 22 BGCs per genome, *Sympodiorosea* have experienced a major loss in biosynthetic potential, likely resulting from a transition from a free-living to a symbiotic/parasitic lifestyle. We hypothesize that the remaining secretome is retained out of necessity for their lifestyle within fungal gardens.

Interestingly, we identified groups of GCFs that correspond to the three described metabolites produced by *Escovopsis weberi*: Melinacidin IV (Fig. S8), shearinine D (Fig. S9) and emodin (Fig. S10) (Heine et al., 2018). In the case of emodin, we identified distinct clustering of the emodin GCFs between the large, derived clade and the clade that contains NGL169, NGL155, NGL214, and NGL159, which may reflect divergent selective pressures acting across the *Sympodiorosea* strains in our study. Considerably more work, however, is needed to capture the depth and breadth of the metabolome in these antagonists. Altogether, the majority of BGCs had no known analog and represent an interesting avenue of discovery. Future work should characterize the specialized metabolism of *Sympodiorosea* to understand how genomic differences in the secretome are manifested. This would help to close the gap in our understanding of the mechanisms of microbial interaction in the fungus-growing ant system.

### Broad patterns of selection in *Sympodiorosea*

Genome streamlining captures the general process by which host-associated organisms lose genes related to a free-living lifestyle (Fijarczyk et al., 2024; Katinka et al., 2001; Moran, 2002; Pombert et al., 2013). This process acts upon antagonistic symbionts associated with fungus-growing ant agriculture and is implicated as the main driver of genomic diversification for *Escovopsis*, *Luteomyces*, and *Sympodiorosea* (Berasategui et al., 2024; de Man et al., 2016; Gotting et al., 2022).

Genome streamlining can work synergistically with purifying selection to reduce functional diversity. Selection of this kind is common in symbionts, where they are experiencing selective pressure from both the hosts to circumvent defenses and competition with the local microbial community (Berasategui et al., 2024; de Man et al., 2016; Fijarczyk et al., 2024; Moran, 2002). In line with other studies of genomic selection in pathogenic fungi, we found genomic variation in *Sympodiorosea* is constrained by purifying selection (Kobmoo et al., 2018; Silva et al., 2015). Combined with the nearly identical ANI between strains, our genomes are consistent with patterns of reduced genomic diversity resultant from a symbiotic lifestyle.

With their evolutionary success largely predicated on streamlining essential processes, symbiont genomes can incur diversity through ‘hotspots’ of evolution, where nonsynonymous mutations accumulate; these genomic regions often contain genes implicated in virulence and defense. We identified a significant difference in selection between the ‘background’ genomic content compared to metabolic genes including BGCs, CAZymes, proteases, and lipases. dN/dS values derived from BGCs were also significantly different than either CAZymes or proteases, suggesting that the *Sympodiorosea* genome evolves in a multi-speed or multi-compartment system (Barelli et al., 2016; Frantzeskakis et al., 2018). Differences between BGCs and other gene groups may indicate that BGCs undergo diversifying selection leading to the maintenance of functional diversity. Ultimately, our findings are consistent with the hypothesis that changes in BGCs are a principal driver of evolution in *Sympodiorosea*.

### Genes with elevated dN/dS

We identified 12 genes with elevated dN/dS values that are implicated in fungal virulence (Fig. S11). dN/dS captures the ratio of nonsynonymous vs. synonymous mutations that accrue in a particular gene. This measure assumes that synonymous mutations in a protein-coding gene will produce a functionally similar protein, whereas nonsynonymous mutations are expected to produce variation that selection can act upon. Nonsynonymous mutations are generally deleterious to the organism. Subsequently, elevated dN/dS indicates an advantage resulting from diversifying selection, which is pertinent to evolution of novel defenses, virulence factors, and for escaping host recognition. Three genes with elevated dN/dS were components of BGCs identified by antiSMASH. Ortho981 had the highest dN/dS value of 0.699. Interestingly this gene encodes an integral membrane protein related to peroxisome maintenance. Peroxisomes are functionally diverse in filamentous fungi and are often implicated in the production of antibiotics and other secondary metabolites (van der Klei & Veenhuis, 2013).

Ortho5377, another gene with elevated dN/dS, likely encodes a telomeric enzyme. In fungi, BGCs are often located in recombinant, sub-telomeric regions of chromosomes where high-repeat nucleotide content and transposable elements often reside. In these regions, genomic stability is lower, and, subsequently, the likelihood of BGCs acquiring variation is higher (Palmer & Keller, 2010; Zhang et al., 2024). Altered telomeric enzymes present an interesting path of evolution for *Sympodiorosea*, particularly as these enzymes may provide an evolutionary ‘playground’ for beneficial mutations to accrue in functionally important regions of the genome.

F-box proteins were also implicated as drivers of genomic diversity in *Sympodiorosea*. These proteins function as major regulators of basic cellular functions as well as highly specific activities, like host-sensing and nutrient acquisition (Jonkers & Rep, 2009; Karahoda et al., 2022; Liu & Xue, 2011). These proteins are often modulated and degraded with ubiquitin-related metabolism, suggesting that transcriptional differences are important in differentiating these fungi from one another. It is thus interesting that we identified multiple instances of F-box protein-encoding genes with elevated rates of selection, which may reflect evolutionary pressure imposed by cultivar presence and chemistry. Also potentially related to cultivar-induced selection, two orthologs of interest likely encode functions related to carbohydrate metabolism. Orth5698 is a CAZyme from the GH3 family primarily responsible for the degradation of cellobiose into glucose. As *Cyphomyrmex* spp. and other lower attine ants provision their gardens with organic detritus, it is possible that these fungi genetically track the most abundant carbon sources in the garden.

Lastly, an extracellular metalloprotease, Ortho5813, was also under elevated rates of selection. Metalloproteases contribute essential functions related to degradation of host tissues, nutrient acquisition and evasion of host defenses. Whether these functions are pertinent in relation to interactions with the cultivars, the ants or other members of the garden community remains an interesting path of investigation, particularly as it pertains to the principal drivers of evolution for *Sympodiorosea*.

### A broader lifestyle than presumed?

*Sympodiorosea* have long been considered as parasites that specialize on attacking and consuming the fungal cultivars grown by lower fungus-growing ant species (Gerardo et al., 2004, 2006; Montoya et al., 2021). We found signatures within the genome consistent with this perspective but also some evidence that *Sympodiorosea* may consume non-fungal tissues. Specifically, in assessing putative substrates of carbohydrate active enzymes encoded in the sequenced genomes, we found strong evidence for utilization of the most abundant polysaccharides in fungi, but we also found evidence for utilization of lignin, starch, and xylem, which are specific to plants. Lignin and xylem are recalcitrant, plant specific polymers critical to cellular structure and nutrient transport. The presence of these enzymes-encoding genes may result from a more complex lifestyle in association with plants, or they may serve an ecological role in the garden by degrading carbohydrates that are otherwise inaccessible. Importantly, when we assessed CAZyme profiles within the context, we found that clustering is likely shaped by shared evolutionary history indicating that these genes evolve neutrally or through intense purifying selection.

It is plausible that *Sympodiorosea* retain the ability to degrade plant matter as a remnant of a past free-living or endophytic lifestyle or that they retain the metabolic capacity to degrade the byproducts produced by cultivars as they degrade plant substrates. This is similar to the parasitic fungus, *Ophiocordyceps australis*. These fungi live a ‘double-life,’ where the fungus can activate specific enzymes in response to an ant or plant infection (de Menezes et al., 2023). While *Sympodiorosea* have never been found to infect plants, it is possible that their introduction to ant gardens coincides with the introduction of fresh material to the garden. It is also plausible that *Sympodiorosea* are tracking the most abundant nutrients in the garden, requiring some degradation of foraged plant matter. With an abundance of fungal derived carbohydrates available, we find the former scenario to be most plausible. Future work should experimentally validate the ability of *Sympodiorosea* to utilize alternative carbohydrates.

While CAZyme diversity tracks phylogenetic diversity, we found much weaker clustering when assessing the serine protease profiles across *Hypocreales* fungi, suggesting that these genes may be important for lifestyle specialization. In this context, *Sympodiorosea* were most similar to *E. weberi*, a specialized mycoparasite of higher attine agriculture, and the free-living saprotroph *Acremonium chrysogenum*. Mycoparasites from *Trichoderma* and plant pathogens from *Ustilaginoidea virens* and *Fusarium graminearum* comprised the nearest cluster to that containing *Sympodiorosea*. Discordance between CAZyme and serine protease genetic variation invites future study of how serine proteases contribute to functional variation in *Sympodiorosea* and the family *Hypocreales* more broadly.

## Conclusion

We sought to understand how selection shapes genomic diversity related to secondary metabolism, with the goal of elucidating potential mechanisms of evolution of these fungi in response to association with the fungus-growing ant symbiosis. Our analysis revealed considerable phenotypic diversity derived from modest genomic diversity in a small pool of samples. *Sympodiorosea* genomic evolution appears to be largely shaped by purifying selection. Exceptions include select genes involved in virulence and membrane protein synthesis, telomeric modification and F-box proteins. Relative to the genome, these genes exhibit diversifying selection, indicating potential pathways for these fungi to specialize to the agricultural lineage of their origin, or perhaps to some other biotic factor to which we are not aware. Combined with our results demonstrating that presence/absence of BGCs serves as a phylogenetic trait, we posit that gain/loss of metabolic genes serves as a putative mechanism for genomic diversification and subsequently, coevolution, between antagonists and the fungal gardens.

Future research should explore the mechanistic basis of evolution through functional characterization of genes of interest and eventual screening of mutants against cultivars to identify altered biological processes. Additionally, the identification of metabolites that modulate interactions between cultivars and *Sympodiorosea* could provide a considerable step towards uncovering the mechanistic basis of fungal evolution in the fungus-growing ant system.

## Data availability

Fungal strains are available upon request. File ‘Supplementary Methods and Figures’ contains detailed descriptions of reagents not included in main text. File ‘Supplementary Tables’ contains sample metadata, assembly statistics, and relevant data for comparative genomics. Sequence data are available at GenBank and the accession numbers are listed in File ‘Supplementary Table S1 and S2’.

## Acknowledgements

We are thankful to Cameron Currie and Ulrich Mueller for providing samples. We also thank the Panamanian Ministerio de Ambiente for collection and export permits. We thank James (Gabe) DuBose for assistance with python scripts related to the selection analysis and Sandy Lin for her assistance in conducting bioassay experiments. Members of the Gerardo and de Roode labs provided valuable feedback.

## Author Contributions

J.R., A.B., N.M. G., and T.D.R. conceived the study. N.M.G., Y.C and H.F.M. collected samples. J.R., A.B., and Q.V.M. performed DNA extractions. J.R., A.B., and T.D.R. assembled genomes. J.R., A.B., Q.V.M., and A.R. performed analyses. J.R. wrote the manuscript. All authors provided valuable comments on the manuscript.

